# Inverse modeling unveils governing law of mechano-chemical dynamics of epithelial migration

**DOI:** 10.1101/2025.09.17.676715

**Authors:** Yuto Kikuchi, Yoshifumi Asakua, Kazuhiro Aoki, Yohei Kondo, Honda Naoki

**Author notes:** Corresponding author(s). E-mail(s): Yohei Kondo Honda Naoki.

## Abstract

Collective cell migration is fundamental to tissue homeostasis and underlies biological processes such as wound healing and cancer invasion. Previous work has proposed governing equations to describe how chemical and mechanical inputs regulate these movements, but the quantitative validity of such models remains to be thoroughly assessed. Here, we developed a machine-learning framework that infers the governing equation from live-cell imaging data. Applied to epithelial sheet migration driven by MAPK/ERK, our approach quantitatively predicted single-cell movement from local chemical and mechanical cues. Examination of the learned equations further indicated that cells process environmental signals by computing their spatiotemporal derivatives. Moreover, when applied to individual cells, our framework revealed cell-cell heterogeneity in the underlying migratory rules. Our framework offers a powerful tool for predictive modeling of multicellular dynamics in both physiological and pathological settings.

## 1 Introduction

Collective cell migration is a fundamental driver of tissue homeostasis and underpins a variety of biological processes, including wound healing and cancer invasion[1–6]. These coordinated movements are orchestrated by intricate biochemical networks, whose disruption is closely associated with pathogenesis[7, 8]. Recent advances in live-cell imaging have enabled the comprehensive tracking of the motion of every cell within a tissue, while simultaneously visualizing intracellular signaling dynamics. This wealth of spatiotemporal data provides unprecedented opportunities to elucidate the mechanistic principles driving collective cell behaviors, a long-sought goal in biological research. To realize this ambition, here we developed a biophysics-tailored machine learning framework that bridges single-cell dynamics and tissue-scale coordination, providing a data-driven pathway to uncover the rules underlying collective cell migration.

Earlier experimental work has pinpointed the mitogen-activated protein kinase (MAPK) extracellular signal-regulated kinase (ERK) as a master regulator of collective migration[9–11]. ERK is a serine/threonine kinase best known for integrating mitogenic cues to control proliferation, differentiation and oncogenic transformation[12–15], yet it has also emerged as a key node linking biochemical signalling to emergent tissue-scale motion. ERK-dependent control of collective behaviour is remarkably conserved across vertebrates, having been observed in the zebrafish and mouse[16, 17]. Among in vitro systems, wound-healing assays in Madin-Darby canine kidney (MDCK) epithelial monolayers have proved uniquely tractable for mechanistic dissection. In this model, ERK activity originates at the wound edge and propagates across the cultured epithelia, whereas cells migrate toward the lesion, suggesting that motility is directed opposite to the travelling ERK wavefront. Using Förster resonance energy transfer (FRET)-based biosensors, we previously captured the spatiotemporal choreography of ERK activity waves and concomitant cell trajectories during MDCK wound closure[21, 22]. Furthermore, by optogenetically imposing synthetic ERK propagation within the monolayer, we demonstrated that the kinase wave itself was sufficient to steer collective migration in situ, underscoring ERK’s causal role in coordinating tissue-scale movement.

The growing body of high-resolution spatiotemporal data on collective cell migration has led to the development of sophisticated mathematical models[21, 23–25]. One of the key remaining challenges is to elucidate the causal link between single-cell motion control and emergent tissue-scale dynamics. From this perspective, we developed a hierarchical modeling approach that translates a particle-based cell-level model into a continuum tissue-level framework[26]. This study not only reproduced tissue motion driven by ERK activity waves but also proposed a plausible mechanism for single-cell motion control during epithelial wound healing.

A qualitative match between simulation outputs and observational data is insufficient to confirm that the model faithfully captures physiological cellular behavior. Moreover, the predictive capability of such mathematical models at the level of individual cell trajectories has yet to be rigorously assessed. These limitations motivate data-driven inverse approaches, wherein time-series data derived from actual cell movements are used to infer the governing mechano-chemical laws underlying their behavior. Such inverse analyses enable the extraction of biologically grounded control principles directly from phenomena—an advance that is not achievable through forward modeling alone.

We presented an inverse analysis framework that uncovers how cells integrate biochemical and mechanical signals to control their movement. Previous mathematical models in this field have often been guided by intuition from researchers and have typically assumed that cells behave as instantaneous reflexes to external cues. In reality, however, cells transform incoming biochemical and mechanical signals through complex intracellular processes before making directional movement decisions. This complexity has largely been overlooked in conventional models. To address this gap, we developed a temporally extended model combined with a biophysics-tailored machine-learning approach. By applying our framework to time-series imaging of signaling activity and cell trajectories, we directly inferred how changes in chemical cues and tissue deformation drove the motion of individual cells.

## 2 Results

This study aimed to uncover how cells translate MAPK/ERK activity and mechanical environments into collective behavior by learning governing principles from time-series data. Specifically, we derived equations linking ERK activity waves with coordinated cell migration during wound healing (Fig. 1A, Supplementary Figure S1 and Supplementary Movie S1). To this end, we introduced a data-driven inverse modeling framework that not only estimated model parameters but also predicted single-cell trajectories in imaging data (Fig. 1B).

**Fig. 1.**
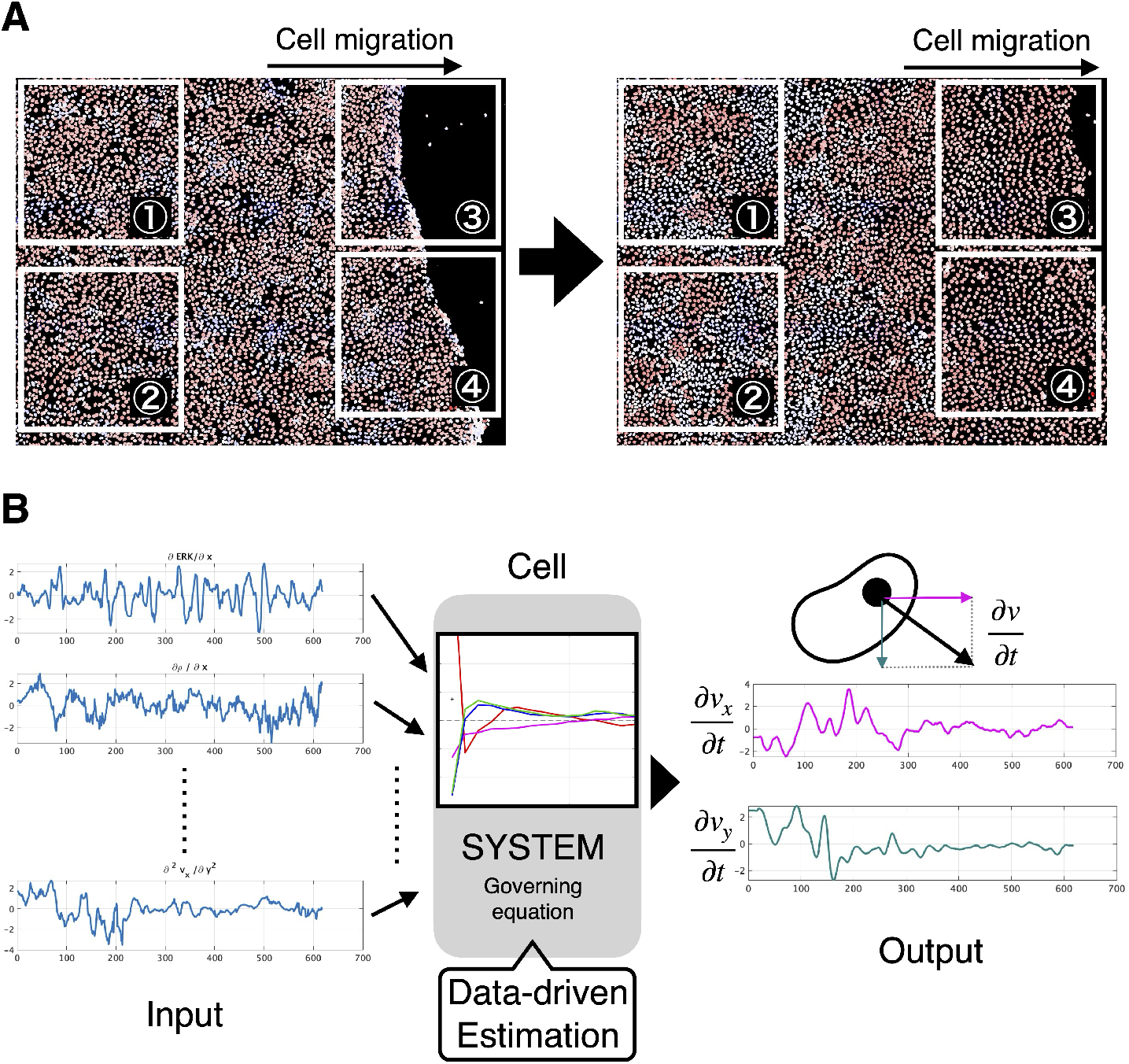
Snapshot of imaging data and framework overview. (A) Snapshots of imaging data of MDCK epithelial cells used for analysis (left: initial state / right: cells spreading into the wounded region). Red indicates high ERK activity, while blue indicates low ERK activity. Over time, cells spread into the wound area. The numbered areas represent the clipped regions used for analysis: 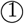 Training data of interior cells, 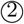 Test data of interior cells, 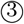 Training data of front cells, and 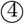 Test data of front cells (See Supplementary Movie S1 and Supplementary Figure S1 for details). (B) Framework overview. Collective cell migration is assumed to be determined by processing multiple input data through an internal system. The aim of this study is to establish a data-driven framework to infer the governing equations, which represent this internal system of the cells.

As the target variable for model prediction, we quantified the acceleration of each cell at every time point based on cell tracking (Fig. 2A). Since we used two-dimensional imaging data, acceleration was decomposed and quantified along two orthogonal directions: vertical and parallel to the wound-healing axis. The input variables to model were from ERK activity and mechanical context. They were also quantified for the same individual cells.

**Fig. 2.**
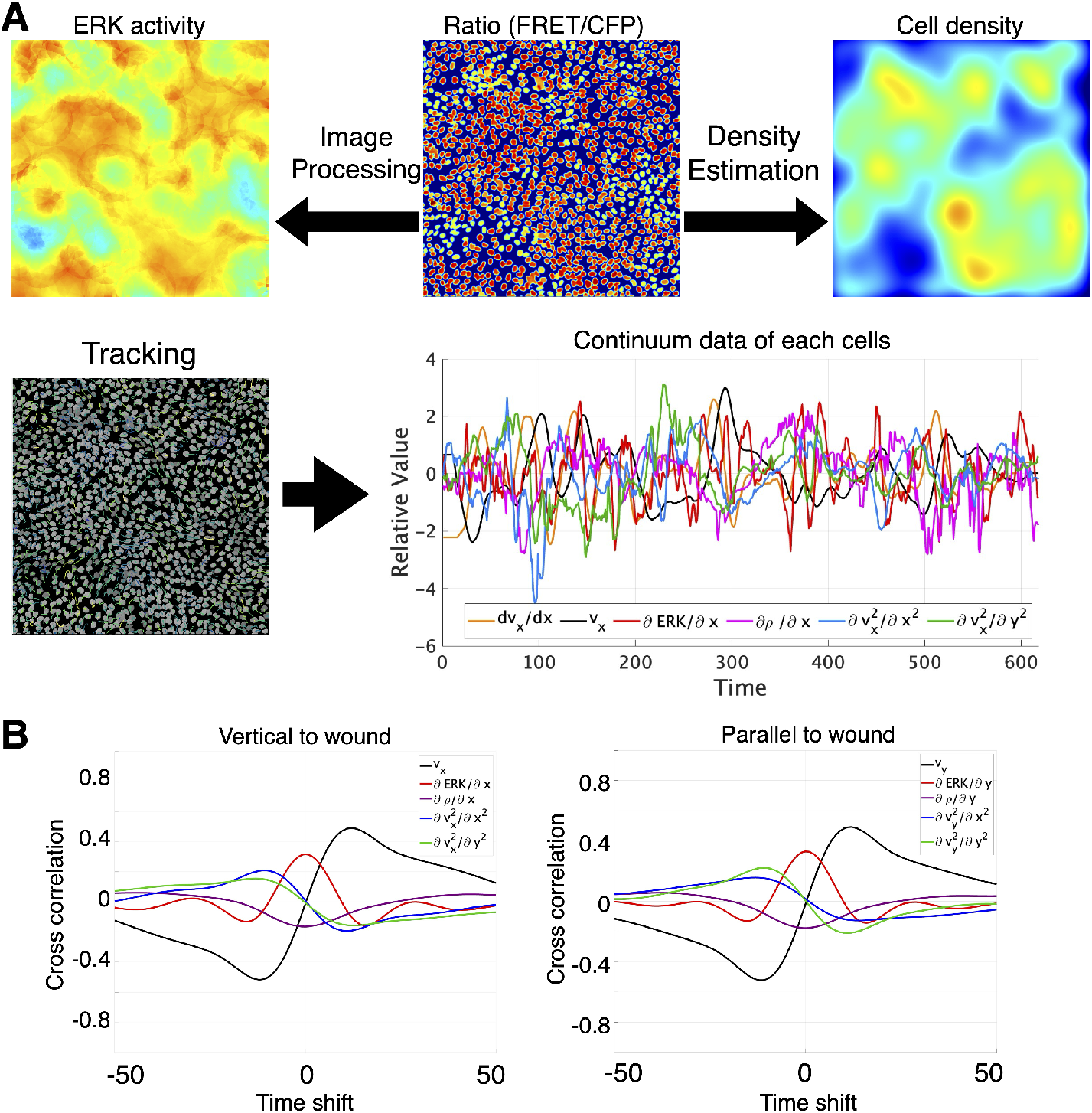
Live imaging of ERK-driven collective migration and extraction of single-cell time-series data. (A)Quantification workflow for the imaging data. Automated cell detection and tracking yield the 2-D position and ERK activity of every cell in each frame. Cell density is computed with kernel density estimation (KDE). Temporal and spatial derivatives are obtained by fitting second-order polynomials to spatio-temporal distributions of ERK activity, cell density, etc. This analysis provides cell acceleration, the spatial gradient of ERK activity, the spatial gradient of cell density, and the second spatial derivatives of cell velocity. The horizontal axis represents elapsed time in frames; images were acquired every 2 min. (B) Mean pairwise cross-correlations of the input features across cells. Correlations were computed for time lags from -100 to +100 min in both the vertical and parallel directions relative to the wound edge. Black, velocity; red, spatial gradient of ERK activity; purple, spatial gradient of cell density; blue, second spatial derivative of velocity along the vertical axis; green, second spatial derivative along the parallel axis.

To predict cell acceleration, we extended our previous framework and proposed the following model:

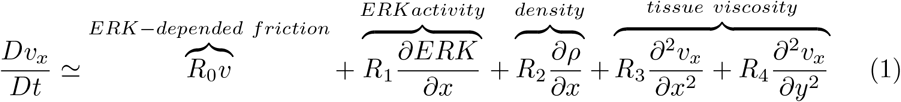

where the coefficients *R*_0_ − *R*_4_ were inferred from the data. This formulation implied that cell acceleration could be approximated as a linear combination of mechanical parameters and ERK activity.

In modeling realistic cell migration, it was important to recognize that the model we constructed might not fully reproduce the experimental data. To enhance the flexibility of the model, we therefore allowed the cell response to depend not only on the current input but also on its recent input history. We referred to the temporally extended coefficients that described this history-dependent response as the response functions.

### Prediction of acceleration and response function

The model used five input features: (i) cell velocity, (ii) the spatial gradient of ERK activity, (iii) the spatial gradient of cell density, and (iv) and (v) the second spatial derivatives of velocity along the x and y axes, respectively. Correlation analysis revealed only weak associations. Cell velocity showed a slight positive correlation with ERK activity and slight negative correlations with its own second spatial derivatives in both directions (Supplementary Fig. 2). Because these pairwise correlations were small, each feature was expected to provide largely independent information to the model. Next, we computed cross correlations between the input features across time lags from 100 to -100 minutes. To assess whether anisotropy existed with respect to the wound edge, we computed cross correlations in the vertical and parallel directions separately (Fig. 2B). This analysis confirmed that the cropped data exhibited no pronounced anisotropy.

Fig. 3A and 3B showed a quantitative agreement between the measured accelerations and those predicted by the inferred model. The cell population was randomly divided into training and test sets, and the test cell with the smallest prediction error was presented. We then compared the shapes of the response functions obtained for the vertical and parallel directions relative to the wound edge (Fig. 3C, 3D). Although a modest discrepancy was detected in the second spatial derivatives, no marked anisotropy emerged. For both directions the response functions displayed differentiator-like behaviour for every feature, implying that cells based their movement on local rates of change rather than on instantaneous feature values alone. In particular, the differentiator response to the ERK gradient may reflect detection of propagating ERK activity waves. We combined the vertical and parallel models to predict the vectorial accelerations (Fig. 3E, 3F), and then we obtained a generally good match, yet several cells remained poorly predicted. Taken together, these results showed that the governing laws inferred from the time-series data accurately explained most cell movements; however, a small fraction of cells—scattered among otherwise well-predicted neighbours—seemed to obey different rules that the current model did not yet capture.

**Fig. 3.**
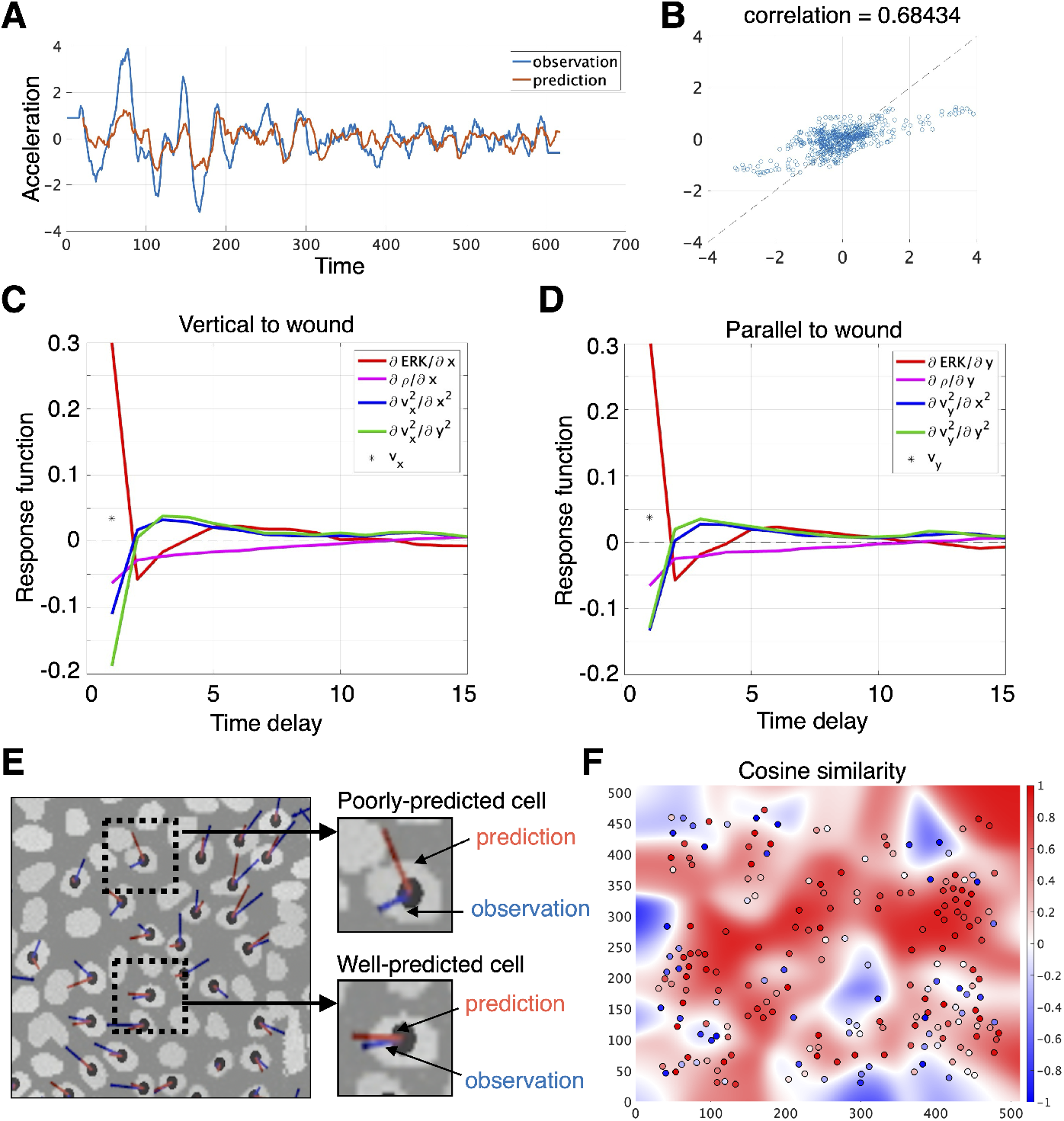
Prediction of single-cell acceleration and inference of response function. (A)Time course of acceleration for a representative cell. Blue, measured values; orange, model predictions. (B) Scatter plot of the same cell’s measured versus predicted accelerations. (C,D) Response functions learned for predicting cell accelerations vertical and parallel to the wound edge.For every input feature except velocity, time lags up to 20 frames (40 min) were considered; only the first 15 frames are displayed here. Black points indicate velocity; red, the spatial derivative of ERK activity; purple, the spatial derivative of cell density; blue, the second spatial derivative along the vertical axis; and green, the second spatial derivative along the parallel axis. (E) Vector-field comparison of observed (blue) and predicted (red) acceleration across the field of view. Vector length indicates magnitude, and orientation indicates direction. Most cells are well captured by the model, although a few poorly predicted cells are also present. (F) Plot and heatmap showing the cosine similarity between observed and predicted acceleration vectors.

### Evaluation of response function properties of cellular units

To elucidate the above-mentioned “rule heterogeneity,” we estimated a distinct response function for every single cell in the test population and assessed the pairwise similarity between cells (Fig. 4). Fig. 4A presented the cosine-similarity matrix of these cell-specific response functions. In the heat-map axes, cells were sorted by how accurately their accelerations were predicted by the population-level response function derived from the training data, arranged from best to worst. Test cells that were well predicted by the training-derived response function exhibited high mutual similarity, as did those that were poorly predicted. Nevertheless, the cell-specific response functions did not form discrete clusters; rather, they lay along a continuous spectrum (Fig. 4A and Supplementary Fig. 4A). We subjected the individual response functions of all test cells to principal-component analysis (PCA) and confirmed the continuous nature (Fig. 4B and Supplementary Fig. 4B). To further characterize the rule heterogeneity, we calculated response functions using two 20-cell subsets: one containing cells that the population-level model predicted accurately and another containing cells it predicted poorly. Fig. 4C and D (and Supplementary Fig. 4C,D) showed the response functions for the vertical-to-wound accelerations, for the well-predicted and poorly-predicted groups, respectively. In the well-predicted group (Fig. 4C), we again observed a biphasic response to the ERK gradient, echoing the behaviour of the training population. Moreover, the viscous drag exerted a positive influence on acceleration after a slight temporal delay, representing the temporal structure of the chemical and mechanical inputs. On the other hand, in the poorly-predicted group (Fig. 4D), the response functions remained qualitatively similar but were generally flatter and of lower amplitude. This profile implied that these cells either lacked a tight coupling between the signals and acceleration or operated under a highly stochastic motility regime. Yet, the high cosine similarity observed within this group indicated that its members still followed a common kinetic rule.

**Fig. 4.**
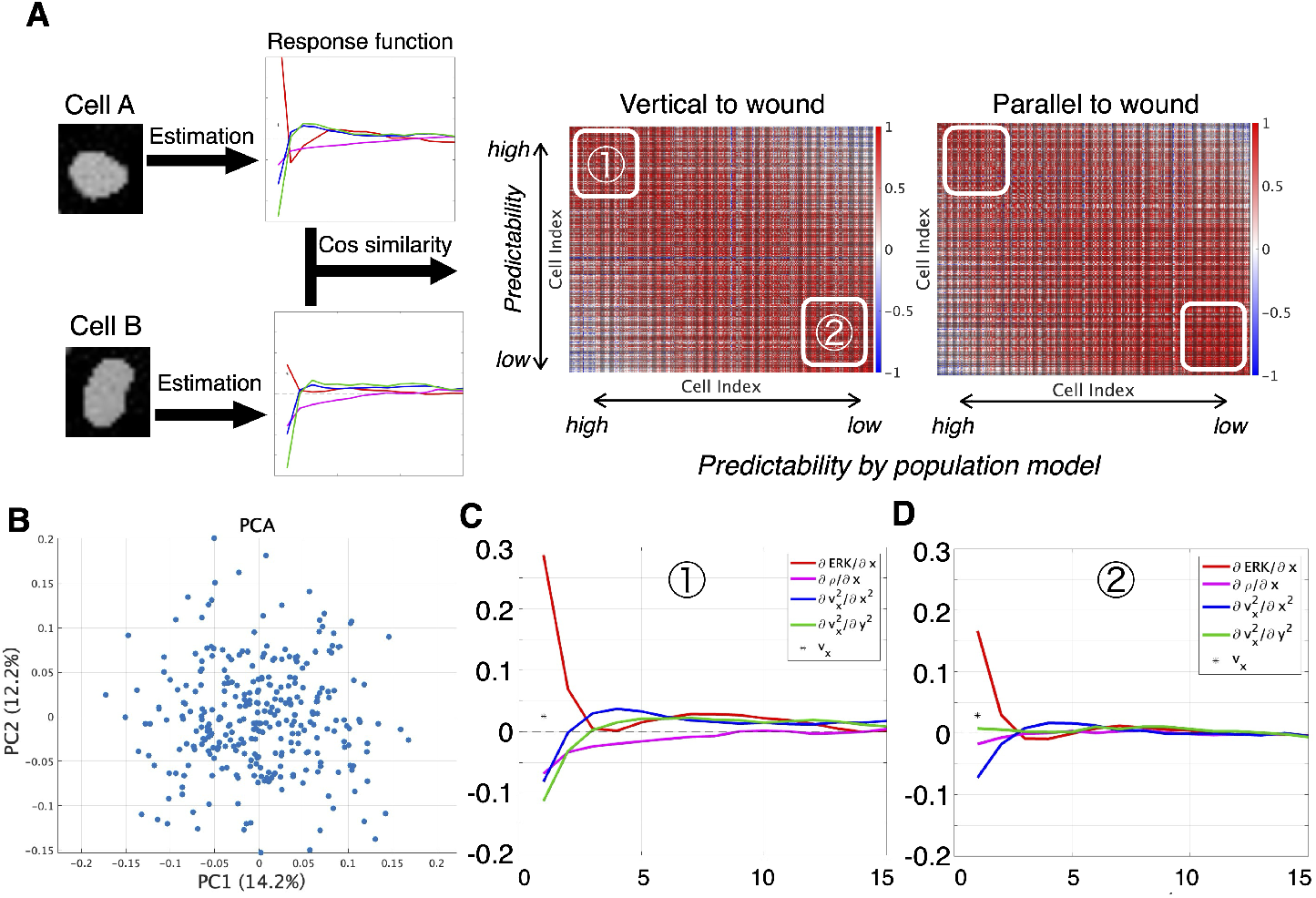
Analysis of cell-cell heterogeneity of response functions. (A) Heat map of pair-wise cosine similarity among response functions estimated from all 304 test-set cells individually. Columns and rows are ordered from left to right (top to bottom) by the prediction performance by the population-level response function from the training-set. Thus, well-predicted cells appear toward the upper left. (B) PCA of the test-cell response functions. Each point represents one cell, summarizing variability in the response patterns. (C) Mean vertical-to-wound response functions for the 20 cells (7 % of the test set) whose accelerations were predicted most accurately by the training-set population-level model. (D) Mean vertical-to-wound response functions for the 20 cells with the poorest prediction accuracy.

### Response function in part of wound-hearing

Until now, our analysis had centred on cells embedded deep within the tissue, far from the wound margin. As exemplified by “leader cells” that emerged at the advancing edge, we reasoned that cells situated close to the wound might follow distinct motility rules. To test this hypothesis, we applied our inference framework to the cell cohort adjacent to the wound and calculated their response functions (Fig. 5A,B). During wound closure, an ERK-activity wave propagated away from the lesion, creating a strong directional bias that drove cells predominantly along the vertical-to-wound axis. Surprisingly, however, the response functions for vertical and parallel motion were highly similar. Thus, even though the extracellular cues were highly anisotropic, the intrinsic law that governed cell movement remained effectively isotropic. A more detailed look at the response to the spatial gradient of ERK activity exposed a clear divergence between front and interior cells. In front cells, the response function peaked at the current ERK level and collapsed to near zero for all positive time lags (Fig. 5C). Interior cells, by contrast, displayed a differentiator-like profile: the response peaked at zero lag, became negative, and then gradually returned to positive values (Fig. 3C,D). This pattern implied that interior cells compared present ERK levels with their recent history when choosing how to move. Collectively, these findings showed that interior cells relied on changes in past ERK activity, whereas front cells near the wound based their behaviour almost exclusively on instantaneous ERK signals (Fig. 5D).

**Fig. 5.**
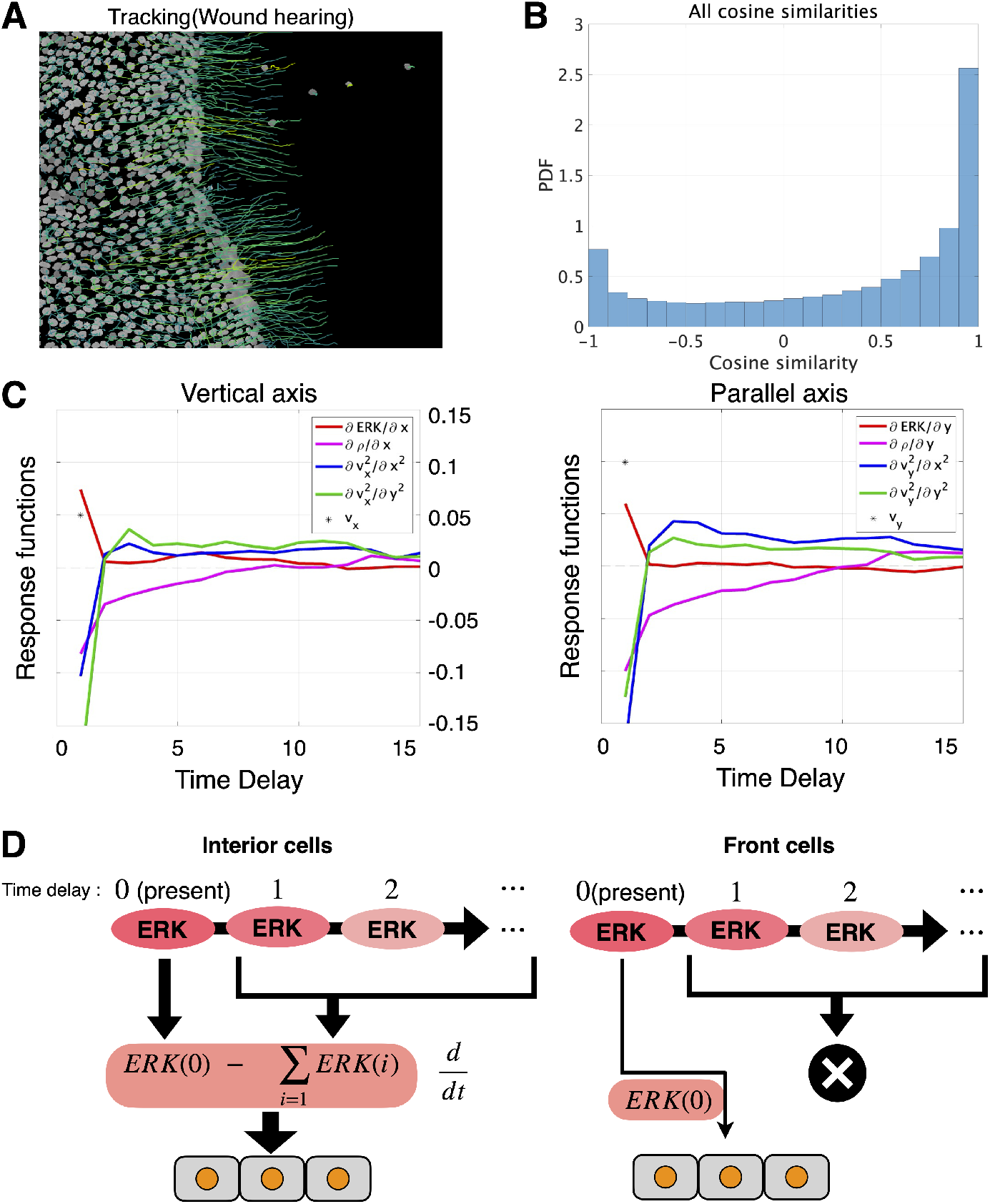
Front-edge cells rely on a motility rule that differs from that of interior cells. (A) Inference of response functions for front cells during wound healing. Left: cell trajectories (100 frames, 200 min) super-imposed on the FRET ratiometry image; the tracks point toward the wound. Isolated cells on the far right and tracks shorter than 100 frames were excluded. (B)Histogram of all cosine similarities. The vertical axis shows the probability density. (C)Response functions for acceleration vertical and parallel to the wound edge, respectively. In front cells, the response pattern to ERK gradient peaks at zero lag and then rapidly decays toward zero. (D) Schematic of ERK-gradient response rules in interior vs. front cells. For interior cells, the response function acting on the ERK gradient is differentiator-like, driving migration according to recent temporal changes rather than absolute levels. In front cells, the response function places virtually all weight at zero lag, indicating that migration depends mainly on the instantaneous ERK gradient.

### Forward analysis of collective cell migration

The data-driven control law we identified behaved like a temporal differentiator and offered clear guidance for refining existing mechanistic models. To illustrate this idea, we revisited the one dimensional particle and spring model [21], in which particles denoted cell positions and springs represented elastic properties. Fig. 6 (and Supplementary Fig. S5) showed a simulation time course of the model. Although the net motion of cells was opposite to the direction of ERK activity waves, our forward simulation revealed a transient movement of cells toward wave propagation upon wave arrival. This behavior was not observed in cultured epithelia. In the original model cell radius changed in response to the instantaneous ERK level. We improved the model so that cell radius instead changed in response to the time derivative of ERK activity as indicated by our data driven response function. This modification removed the artifact of cells moving with the wave (Fig. 6C and Supplementary Fig. S5B).

**Fig. 6.**
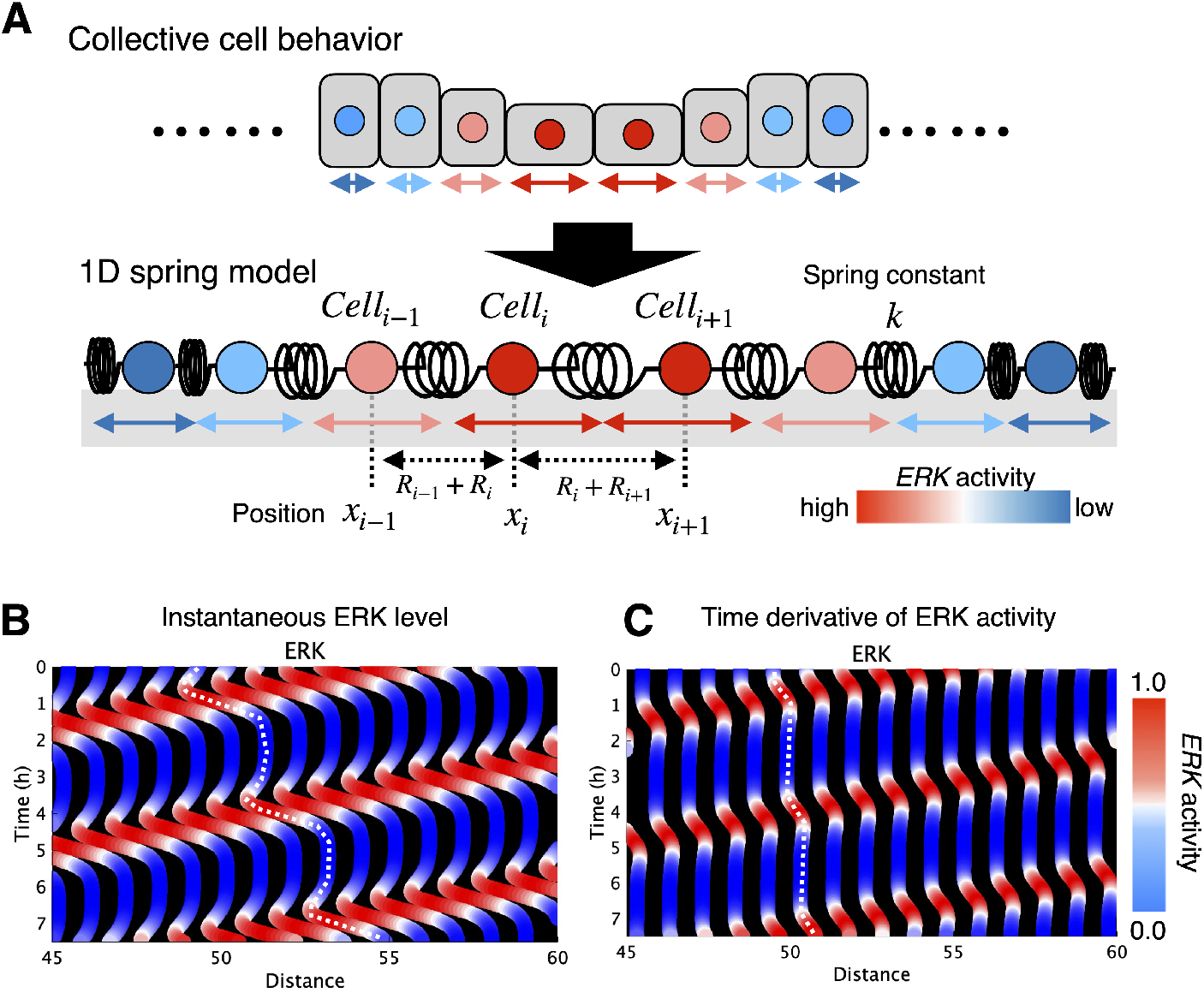
Forward analysis of collective cell migration. (A)Numerical simulation of one dimensional particle and spring model. (B)The instantaneous ERK level(Eq. 3 in previous study). (C)The time derivative of ERK activity. Horizontal axis is cell scan position (Distance, 45-60) and vertical axis is elapsed time since experiment start (Time, 0-7 h). Color denotes the magnitude of each value. These panels visualizes an ERK activity wave propagating from right to left over time. Parameter and setting of (a) are k = 2 (min 2), *µ*_0_ = 10 (min 1), *R*_0_ = 1/2, *α* = 1.5, *β* = 2.5, *σ* = 0.1 (min 1), sweeping velocity of illumination = 0.1 (min 1), width of illuminated area = 30, number of cells = 100. Parameter and setting of (b) are k = 2 (min 2), *µ*_0_ = 10 (min 1), *R*_0_ = 1/2, *α* = 10, *β* = 2.5, *σ* = 0.1 (min 1), sweeping velocity of illumination = 0.1 (min 1), width of illuminated area = 30, number of cells = 100. Where *k* is the spring constant between neighboring cells; *R*_0_ and *µ*_0_ are the basal cell radius and viscosity, respectively, modulated by ERK activity with amplitudes *α*(size) and *β*(viscosity); and *σ*is the ERK decay rate.

## 3 Discussion

In this study, we developed an inverse-analysis framework that uncovered governing principles of cell motility grounded in quantitative biological evidence and leveraged them to predict cell movements with high fidelity. We applied the framework to simultaneously acquired time-series data of ERK activity and cell migration in cultured epithelial sheets obtained through FRET-based live imaging. The analysis revealed that the trajectories of individual cells could be quantitatively predicted from the combined spatiotemporal gradients of mechanical conditions and chemical cues. Because our framework derived control laws directly from experimental time-series data, it provided a broadly applicable, data-driven route for extracting biologically meaningful regulatory rules from observed cellular dynamics.

The derived response functions revealed a distinctive temporal pattern for the spatial gradient of ERK activity. Specifically, the function exhibited a sharp positive peak at zero time lag, followed by a rapid decline that overshot into a transient negative phase. This biphasic shape implied that cells responded to ERK signals in a differentiator-like manner, generating mechanical reactions that were acutely sensitive to the current rate of change rather than to the absolute level of activity. The delayed negative component further indicated that cells recognized not only the initial rise but also the subsequent fall of ERK activity, suggesting an ability to anticipate—or at least account for—the dynamic profile of the ERK gradient arising from wave propagation. By contrast, the response functions for the density gradient and the second spatial derivatives of velocity field were smoother and more sustained, indicating that these mechanical cues acted on a slower timescale and became prominent after the sharp chemical trigger provided by ERK activity. Of note, the density-gradient response likely reflected an elastic counterforce, whereas the velocity-gradient response may capture a viscous one. Together, these findings supported a two-step control scheme in which a rapid chemical stimulus initiated movement, followed by mechanical feedback that complemented and modulated the ensuing migration.

When we compared response-function similarity at the single-cell level, we found that cells that were accurately captured by the population-average model shared similarly shaped response functions—and likewise for cells that the population model predicted poorly. Interestingly, similarity analysis across all cells showed no discrete clusters; rather, the response functions formed a continuous spectrum. In other systems, however, truly discrete clusters might exist, and our approach should have been able to uncover such hidden cell types. In those cases, relying on a single population-average model would have been inadequate, making it important to extend our framework with hierarchical modeling.

At the wound front, we uncovered control rules that differed from those operating in interior cells, most strikingly for the chemical cue represented by the spatial derivative of ERK activity. Wound-front cells lacked the biphasic response to the ERK gradient observed in interior cells and instead exhibited a single positive-peaked response. In other words, these cells did not behave as differentiators that read the rate of change of ERK gradient; they accelerated in proportion to the current ERK level. Consistent with this, sustained ERK activation was frequently observed at wound margins, suggesting that front cells employed motion-control rules tuned to a persistent chemical signal. The density gradient, by contrast, exerted an overall inhibitory influence, suppressing movement toward the high-density interior. Collectively, these results indicated that epithelial collectives deployed distinct information-processing strategies according to local conditions and demonstrated that our framework could identify such heterogeneity in control rules.

Here, we discussed the limitations of our current framework. The regression model we presently employed assumed linearity, leaving potential nonlinear effects largely unexamined. Expanding the model to incorporate nonlinear interactions would have made it possible to capture higher-order crosstalk among additional MAPK family members and mechanosensory pathways, yielding a more comprehensive view of the control architecture governing cell motility. In addition, cell collectives likely adapted to non-stationary environments by dynamically re-weighting chemical and mechanical cues that guide their behavioral choices. The culture-based framework developed in this study did not yet accommodate such flexible signal weighting. A recent machine-learning approach aimed at outdoor analyses of medaka gonadal development addressed this need by inferring adaptive signal weights from environmental cues and physiological states[27]. Incorporating a similar strategy—estimating model-parameter weights directly from environmental factors and cellular time-series data—into our regression pipeline could have enabled us to derive context-dependent control laws in vivo from experimental data.

The machine-learning of response functions not only allowed us to predict cell migration but also informed targeted improvements to an existing mathematical model (Fig 6). Moreover, these data-driven insights furnished a principled foundation for probing the specific molecular pathways that implement the inferred control behaviors, thereby offering a valuable framework for guiding subsequent experimental validation.

This study advanced both the mechanistic understanding of multicellular dynamics and the methodology of data-driven modeling. By inferring governing laws directly from time-series imaging, we quantitatively uncovered a mechanochemical control law that connected single-cell mechanobiology to tissue-scale behavior. Grounded in a general regression framework, the method could be transferred from MDCK monolayers to a broad array of multicellular systems, opened new avenues for quantitative analyses of collective motion in biomedical and engineering contexts. We believed these results laid a foundation for extracting governing principles from complex phenomena that had resisted traditional bottom-up modeling.

## 4 Materials and Methods

### Image analysis

We outlined the image-analysis workflow used to extract single-cell features for training our machine-learning model. ERK activity waves during scratch-wound healing were visualized previously with a Förster resonance energy transfer (FRET) biosensor and time-lapse microscopy[21]. From these movies we extracted cell trajectories with TrackMate, an ImageJ/Fiji plug-in that links detected cells across frames via cost-minimizing assignment[28–30]. The resulting cell coordinates were processed with a second-order Savitzky-Golay filter (window = 11 frames) to obtain first- and second-order temporal derivatives, that is, cell velocities and accelerations, respectively. In addition, we obtained spatial derivatives of the velocity field by least-squares fitting a quadratic polynomial to the velocity vectors of neighboring cells within a radius *r* around each cell.

FRET and CFP channels were first smoothed with a Gaussian kernel (*σ* = 1.0px) to attenuate high-frequency noise. After subtracting background fluorescence, we computed pixel-wise FRET/CFP ratio images for every time point. The ratio value at each cell centroid served as a single-cell measure of ERK activity. Analogous to the velocity analysis, ERK activity values of neighboring cells within a radius *r*(= 10) were least-squares fitted to a quadratic surface, and analytical differentiation of the fitted coefficients yielded the local spatial gradient of ERK activity.

Cell density fields were reconstructed for every frame by kernel density estimation, superimposing a two-dimensional Gaussian kernel (*σ* = 25px) at each cell position. Spatial gradients of the resulting density maps were calculated with the same local quadratic fitting procedure described above.

### Mathematical model

We previously proposed a mathematical model of epithelial cell migration driven by propagation waves of ERK activity[21, 26]:

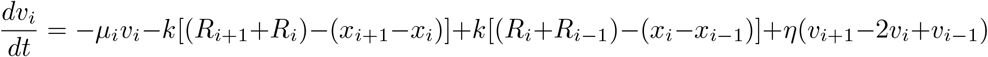

In this model, each cell *i* was characterized by several parameters: its velocity *v*_*i*_, viscosity *η*_*i*_, radius *R*_*i*_, and friction coefficient *µ*_*i*_. The elasticity of the interaction was described by a spring constant *k*, which was assumed to be common to all cells. We subsequently derived a coarse-grained hydrodynamic description of the particle-based model to facilitate analytical analysis. In our previous work[26], we derived a one-dimensional continuum version of the model and heuristically extended it to two dimensions. In the present study, we formally derived the full two-dimensional continuum description directly from the original particle-based model (see Supplementary Text S1 for the derivation).

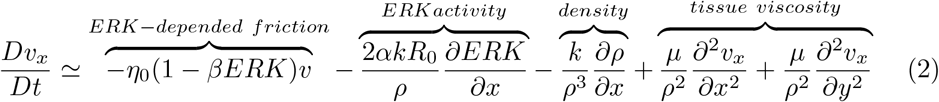

In this model, *v*_*x*_ and *v*_*y*_ represented velocities in the vertical and parallel directions relative to the wound axis, respectively; *η* was the friction coefficient; *β* was a scaling factor for ERK-dependent friction; *ERK* denoted the molecular activity of ERK; *α* was a scaling factor relating ERK activity to cell size; *k* was the elastic (spring) constant; *R*_0_ was the basal cell radius; *ρ* was the density; and *µ* was the viscosity coefficient. This equation indicated that the acceleration of cells could be approximated as a linear combination of mechanical parameters and spatial derivatives of ERK activity. Note that the derivation introduced the term *∂*_*xx*_*v*_*x*_ and *∂*_*yy*_*v*_*x*_, which was absent in the model proposed in our previous work. This term was expected to be effective for capturing the spatiotemporal patterns of cell velocity observed in the data. By substituting the coefficient of each term with *R*_0_–*R*_4_, we obtained Eq.(1) in the main text.

### 1D Spring model

We introduced the mathematical model used in Fig. 6, which was composed of a one-dimensional (1D) array of cells. Since each cell adhered to its neighbors, cells mechanically interacted through repulsive and attractive forces. To represent these mechanical interactions, we assumed a system in which particles were connected by springs. Here, each particle represented the centroid position of a cell, and the springs represented the elastic properties of the cells, including membrane, cytoskeleton, and adhesion. Accordingly, the dynamics of the position of the *i*-th cell *x*_*i*_ were described as follows:

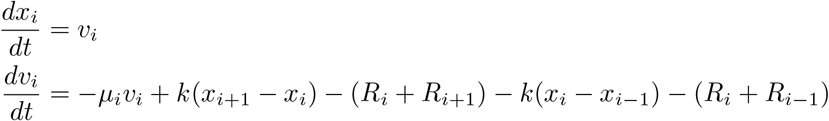

where *v*_*i*_ denoted the velocity of the *i*-th cell, and *k* represented the spring constant of the cell. At the boundaries of the particle array, the end particles were coupled to only one neighboring particle.

In our previous model used in Fig. 6B, *R*_*i*_ and *µ*_*i*_ were modulated by the ERK activity as follows:

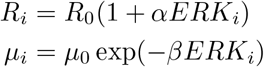

Here, *ERK*_*i*_ denoted the ERK activity of the *i*-th cell, while *R*_0_ and *µ*_0_ represented the basal cell radius and basal viscosity, respectively. Parameters *α* and *β* described the effects of ERK activity on cell size and viscosity, respectively.

On the other hand, in Fig. 6C, we modified the following update rule for the basal cell radius:

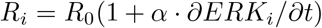

We modeled optogenetically induced ERK activation, which was expressed as:

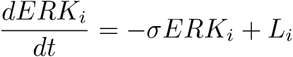

where s and *L*_*i*_ denoted the decay rate of ERK activity and the light intensity applied to the *i*-th cell, respectively. *L*_*i*_ = 1 for illuminated cells while = 0 otherwise. The sweeping velocity of illumination was set to 0.1 (min^*−*1^), and the width of the illuminated area was 30.

### Dataset preparation

We separately cropped non-overlapping regions from the entire field of view of the imaging data, which were used to prepare the training and test sets. To exclude cells that moved into or out of the field of view or appeared through cell division, we limited the analysis to trajectories lasting at least 400 frames. This procedure removed unstable trajectories and extremely short time series that would otherwise have added noise during training. For the same reason, cell trajectories containing missing values were discarded. The resulting training set comprised 398 cell trajectories, each beginning at the first frame and extending for at least 400 frames. The test set contained 303 trajectories processed in the same way. No data augmentation was applied in this study.

### Machine learning

We extended the continuum model (Equation 2) to incorporate the time-series profiles of the mechano-chemical inputs, thereby increasing its quantitative ability to fit real-world data. Past information was incorporated by constructing, for each explanatory variable, a temporal convolution matrix. Specifically, for the response variable, cell acceleration, we assumed the following convolutional model:

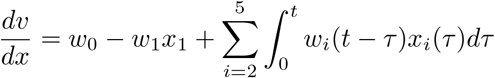

In the regression model, the explanatory variables included a bias term *w*_0_, the velocity component *x*_1_ = *v*_*x*_, the spatial gradient of ERK activity *x*_2_ = *∂ERK/∂x*, the spatial gradient of cell density *x*_3_ = *∂ρ/∂x*, and the second spatial derivatives of velocity, *x*_4_ = *∂*^2^*v*_*x*_*/∂x*^2^ and *x*_5_ = *∂*^2^*v*_*x*_*/∂y*^2^. Here, *τ* denoted the temporal lag (time shift), and *w*_*i*_(*t − τ*) was the weight for the *i*-th predictor at that lag. Because cell acceleration is the time derivative of velocity, allowing an arbitrary temporal function for velocity would have let the model approximate acceleration almost exactly, thereby masking the contributions of the other predictors. For this reason, velocity was included in the model only at the current time point, with no temporal history. After temporal discretization, the convolutional formulation became a multivariate linear regression problem that we fit by ridge (*l*^2^-regularized) regression. The regularization parameter was optimized by five-fold cross-validation, yielding an optimal value of *λ* = 784.760 (Supplementary Fig. 3). The length of the temporal history considered was limited to 20 frames, chosen from the inter-predictor cross-correlation structure observed in the training data (Fig. 2B).

## Code availability

This framework and statistical analysisis in this study developed under MATLAB R2023a. The framework and the statistical analysis code will be distributed through GitHub after publication.

## Data availability

This study was a re-analysis of existing data (Aoki et al., Dev. Cell, 2017). The datasets will be distributed through GitHub after publication.

## Acknowledgements

We are grateful all members of the Honda laboratory for the valuable discussions. This work was supported [Grant Number JPMJSP2132 to Y.Kikuchi] by JST SPRING, Grant-in-Aid for Scientific Research (C) [Grant Number JP25K09578 to Y.Kondo] by JSPS, Moonshot R&D-MILLENNIA Program [Grant Number JPMJMS2024-9 to N.Honda and Y.Kondo] by JST, and the Cooperative Study Program of Exploratory Research Center on Life and Living Systems (ExCELLS) [Program Number 21-102 to N.Honda].

## Author Contributions

N.Honda, K.Aoki and Y.Kondo conceived the project. N.Honda, Y.Asakura developed the model. K.Aoki, N.Honda conducted the experiments. Y.Kikuchi analyzed the data. Y.Kikuchi, Y.Kondo and N.Honda wrote the manuscript with input from all the authors.

## References

[1] K.J. Cheung, E. Gabrielson, Z. Werb, A. J. Ewald: Collective invasion in breast cancer requires a conserved basal epithelial program. Cell 155, 1639–1651 (2013) 10.1016/j.cell.2013.11.029

[2] V. Padmanaban, I. Krol, T. Suhail, B.M. Szczerba, N. Aceto, J.S. Bader, A.J. Wald: E-cadherin is required for metastasis in multiple models of breast cancer. Nature 573, 439–444 (2019) 10.1038/s41586-019-1526-3

[3] N.M. Novikov, J. Gao, A.I. Fokin, N. Rocques, G. Chiappetta, K.D. Rysenkova, D.J. Zea, A. Polesskaya, J. Vihn, R. Guerois, A.M. Gautreau: NHSL3 controls single and collective cell migration through two distinct mechanisms. Nature Communications 16 (2025) 10.1038/s41467-024-55647-3

[4] P. Friedl, D. Gilmour: Collective cell migration in morphogenesis, regeneration and cancer. Nat. Rev. Mol. Cell Biol 10, 445–457 (2009) 10.1038/nrm2720

[5] B. Ladoux, R.M. Méga: Mechanobiology of collective cell behaviours. Nature Reviews Molecular Cell Biology 18, 743–757 (2017) 10.1038/nrm.2017.98

[6] X. Trapet, M.R. Wasserman, T.E. Angelini, E. Millet, D.A. Weitz, J.P. Butler, J.J. Fredberg: Physical forces during collective cell migration. Nature Physics 5, 426–430 (2009) 10.1038/nphys1269

[7] I.V. Lijakovic, M. Milivojevic, A.G. Clark: Collective Cell Migration on Collagen-I Networks: The Impact of Matrix Viscoelasticity. Frontiers in Cell and Developmental Biology 10 (2022) 10.3389/fcell.2022.901026

[8] T.J. Aikin, A.F. Peterson, M.J. Pokrass, H.R. Clark, S. Regot: MAPK activity dynamics regulate non-cell autonomous effects of oncogene expression. eLife (2020) 10.7554/eLife.60541

[9] S. Matsubayashi, M. Ebisuya, S. Honjoh, E. Nishida: ERK Activation Propagates in Epithelial Cell Sheets and Regulates Their Migration during Wound Healing. Current Biology 14, 731–735 (2004) 10.1016/j.cub.2004.03.060

[10] D.L. Nikolic, A.N. Boettiger, D. Bar-Sagi, J.D. Carbeck, S.Y. Shvartsman: The role of boundary conditions in an experimental model of epithelial wound healing. American journal of physiology-cell physiology 8544, 68–75 (2006) 10.1101/2025.03.31.646257

[11] K. Aoki, K. Kumagai, A. Sakurai, N. Komatsu, Y. Fujita, C. Shionyu, M. Matsuda: Visualization of small GTPase activity with fluorescence resonance energy transfer-based biosensors. Molecular cell 52, 529–540 (2013) 10.1016/j.molcel.2013.09.015

[12] L. Chang, M. Karin: Mammalian MAP kinase signalling cascades. Nature 410, 37–40 (2001) 10.1038/35065000

[13] G. Pearson, F. Robinson, T.B. Gibson, B. Xu, M. Karandikar, K. berman, M.H. Cobb: Mitogen-activated protein (MAP) kinase pathways: regulation and physiological functions. Endocrine Reviews 22, 153–183 (2001) 10.1210/edrv.22.2.0428

[14] P.L. Roberts, C.J. Der: Targeting the Raf-MEK-ERK mitogen-activated protein kinase cascade for the treatment of cancer. Oncogene 26, 3291–3310 (2007) 10.1038/sj.onc.1210422

[15] D.G. martinez, L. Roth, T.R. mumford, J. Guan, A. Le, R.C. Doebele, B. Huang, A. and, M.N. Bugaj, T.G. Bivona, L.J. Bugaj: Oncogenic EML4-ALK assemblies suppress growth factor perception and modulate drug tolerance. Nature Communications 15, 9473 (2024) 10.1038/s41467-024-53451-7

[16] T. Hiratsuka, Y. Fujita, N. Honda, K. Aoki, Y. Kamioka, M. Matsuda: Intercellular propagation of extracellular signal-regulated kinase activation revealed by in vivo imaging of mouse skin. eLIFE 4 (2015) 10.7554/eLife.05178

[17] A. De Shimone, M.N. Evanitsky, L. Hayden, B.D. Cox, J. Wang, V.A. Tornini, J. Ou, A. Chao, K.D. Poss, S. Di Talia: Control of osteoblast regeneration by a train of Erk activity waves. Nature 590, 129–133 (2021) 10.1038/s41586-020-03085-8

[18] M. Poujade, E.G. Mongrain, A. Hertzog, J. Jouanneau, P. Chavriert, B. Ladoux, A. Buguin, P. Silberzan: Collective migration of an epithelial monolayer in response to a model wound. Proceedings of the National Academy of Sciences 104, 15988–15993 (2007) 10.1073/pnas.0705062104

[19] R. Farooqui, G. Fenteany: Multiple rows of cells behind an epithelial wound edge extend cryptic lamellipodia to collectively drive cell-sheet movement. Journal of cell science 118, 51–63 (2005) 10.1242/jcs.01577

[20] K. Aoki, Y. Kunigami, A. Sakurai, N. Komatsu, Y. Fujita, C. Shionyu, M. Matsuda: Stochastic ERK activation induced by noise and cell-to-cell propagation regulates cell density-dependent proliferation. Molecular cell 52, 529–540 (2013) 10.1016/j.molcel.2013.09.015

[21] K. Aoki, Y, Kondo, N. Honda, T. Hiratsuka, R.E. ito, M. Matsuda: Propagating Wave of ERK Activation Orients Collective Cell Migration. Development Cell 43, 305–317 (2017) 10.1016/j.devcel.2017.10.016

[22] K. Aoki, M. Matsuda: Visualization of small GTPase activity with fluorescence resonance energy transfer-based biosensors. Nature Protocols 4, 1623–1631 (2009) 10.1038/nprot.2009.175

[23] X. Serra-Picamal, V. Conte, R. Vincent, E. Anon, D.T. Tambe, E. Bazellieres, J.P. Butler, J.J. Fredberg, X. Trepat: Mechanical waves during tissue expansion. Nature Physics 8, 628–634 (2012) 10.1038/NPHYS2355

[24] N. Hino, L. Rossetti, A. Marín-Llauradó, K. Aoki, X. Trepat, M. Matsuda, T. Hirashima: ERK-Mediated Mechanochemical Waves Direct Collective Cell Polarization. Developmental Cell 53, 646–660 (2020) 10.1016/j.devcel.2020.05.011

[25] D. Boocock, T. Hirashima, E. Hannezo: Interplay between Mechanochemical Patterning and Glassy Dynamics in Cellular Monolayers. PRX LIFE 1, 013001 (2023) 10.1103/PRXLife.1.013001

[26] Y. Asakura, Y. Kondo, K. Aoki, N. Honda: Hierarchical modeling of mechanochemical dynamics of epithelial sheets across cells and tissue. scientific reports 11, 4069 (2021) 10.1038/s41598-021-83396-6

[27] T. Itoh, Y. Kondo, T. Nakayama, A. Shinomiya, K. Aoki, T. Yoshimura, N. Honda: Inverse signal importance in real exposome: How do biological systems dynamically prioritize multiple environmental signals? bioRxiv (2025) 10.1101/2025.03.31.646257

[28] J.Y. Tinevez, N. Perry, J. Schindelin, G.M. Hoopes, G.D. Reynolds, E. Laplantine, S.Y. Bednarek, S.L. Shorte, K.W. Eliceiri: TrackMate: An open and extensible platform for single-particle tracking. Methods 115, 80–90 (2017) 10.1016/j.ymeth.2016.09.016

[29] C.A. Schneider, W.S. Rasband, K.W. Eliceiri: NIH Image to ImageJ: 25 years of Image Analysis. Nat Methods 9, 671–675 (2012) 10.1038/nmeth.2089

[30] J. Schindelin, et.al.: Fiji -an Open Source platform for biological image analysis. Nat Methods 9 (2019) 10.1038/nmeth.2019

