## Supplemental information(PDF) for "Inverse modeling unveils governing law of mechano-chemical dynamics of epithelial migration"

### Supplementary Figures

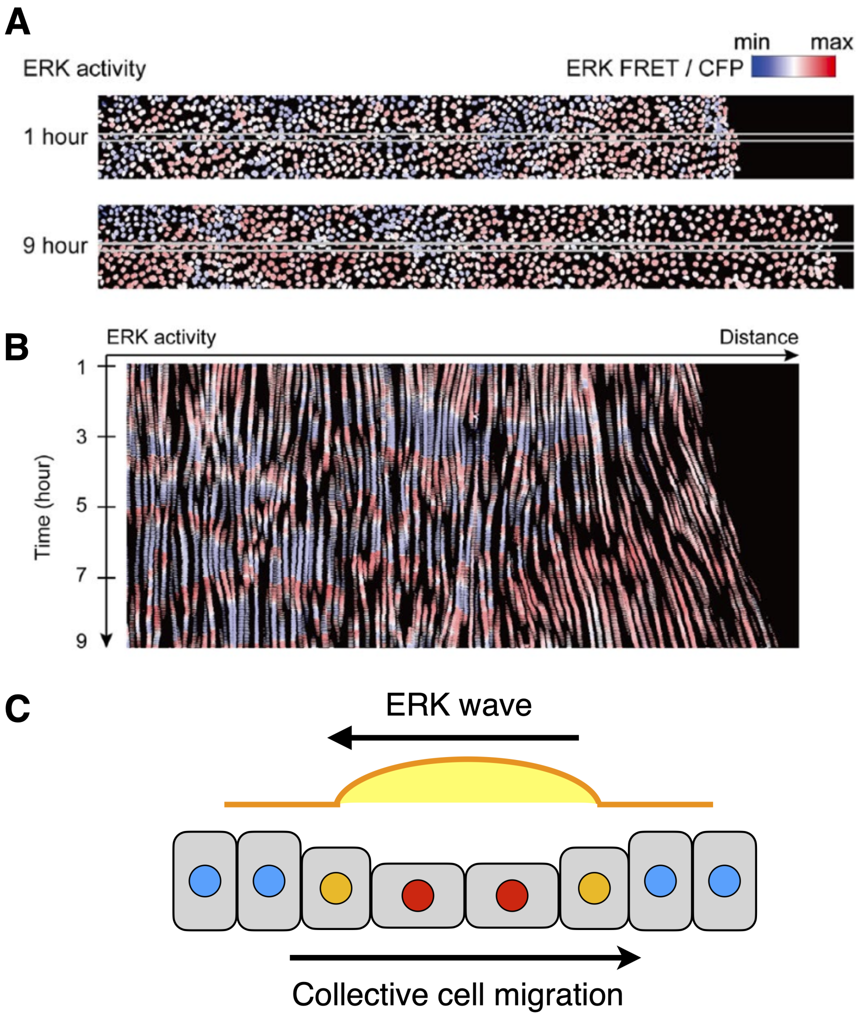


**Supplementary Fig S1** (A) Time-lapse FRET imaging of ERK activity during wound healing in an epithelial MDCK monolayer expressing a nuclear-localized ERK biosensor. Snapshots are shown 1 h (upper) and 9 h (lower) after scratching. Red and blue denote high and low ERK activity, respectively; the cytosol is not visualized because the biosensor is confined to the nucleus. (B) Kymograph of ERK activity in the band region outlined by the white box in (a). (C) Schematic of ERK-mediated collective migration. As an ERK activity wave travels, cells migrate in the opposite direction.

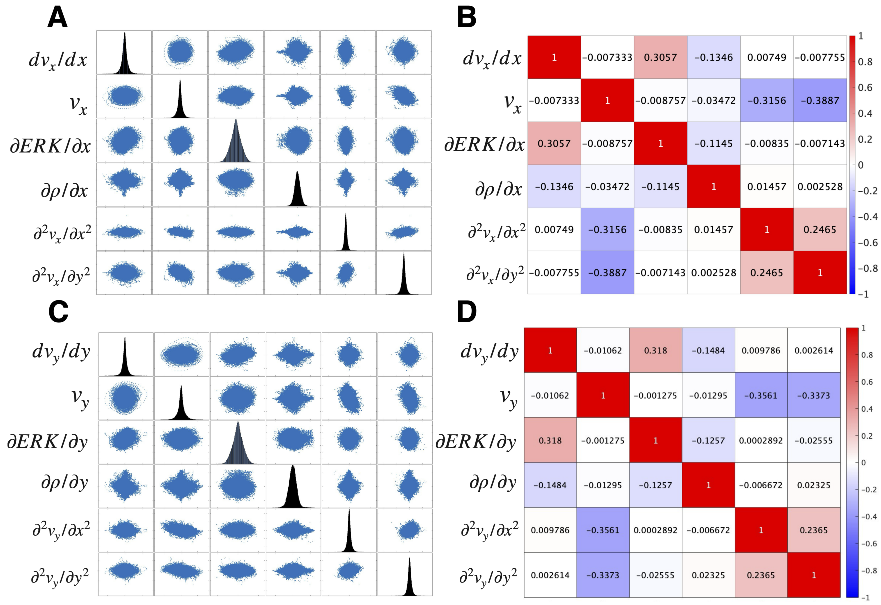


**Supplementary Fig S2** (A) Scatter plot of the input variables supplied to the machine-learning model that predicts cell acceleration vertical to the wound edge. (B)Heat map of the input features shown in (A). A weak positive correlation is observed between cell acceleration and the spatial gradient of ERK activity, whereas cell velocity shows weak negative correlations with both of its corresponding second spatial derivatives. (C) Scatter plot of the input variables for predicting cell acceleration parallel to the wound edge. (D) Heat map of the input features shown in (C). Similar weak correlations are observed, with acceleration positively linked to ERK gradients and velocity negatively linked to its second derivatives.

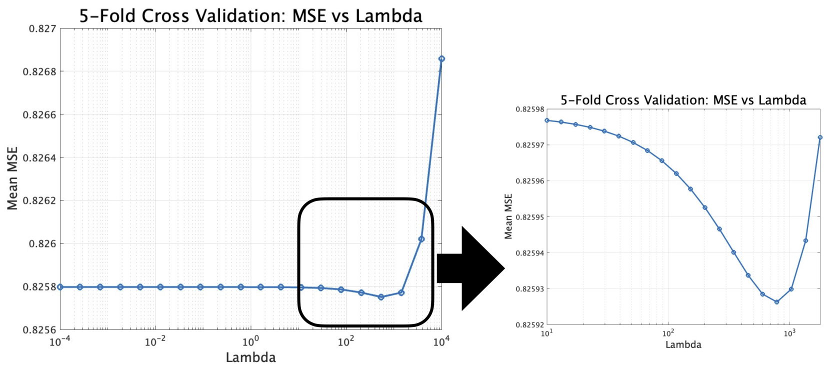


**Supplementary Fig S3** Results of k-fold cross-validation used to optimize the regularization parameter $\lambda$. Only the training set was employed. The left panel reports the cross-validation error over a broad search range ($10^{-4} \leq \lambda \leq 10^{4}$). To locate the minimum, cross-validation was repeated within a narrower neighbourhood ($10^{-1} \leq \lambda \leq 10^{3.75}$), yielding an optimal value of $\lambda = 784.7600$.

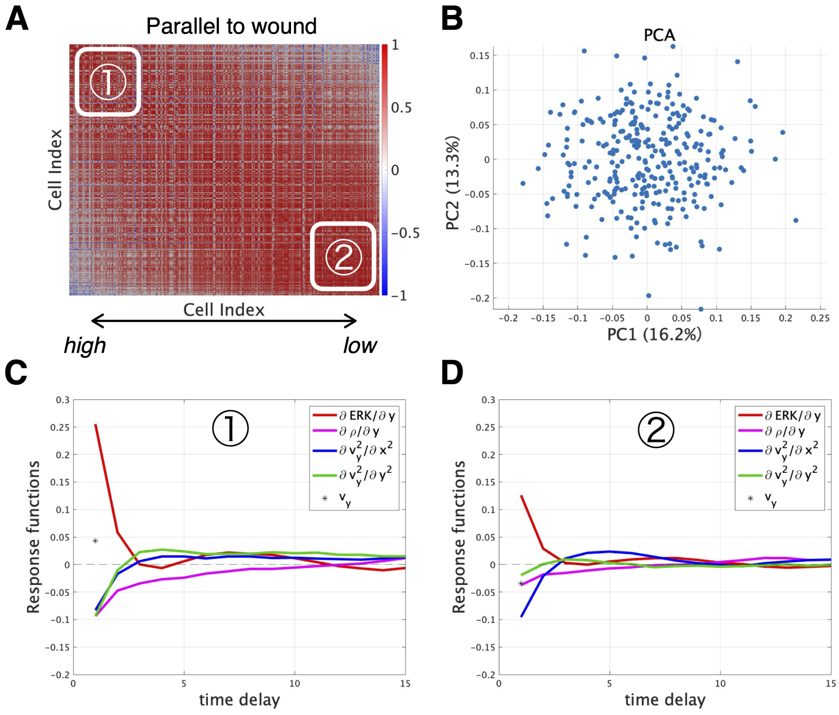


**Supplementary Fig S4** (A) Heat map of pair-wise cosine similarity among response functions estimated from all 304 test-set cells individually. Columns and rows are ordered from left to right (top to bottom) by the prediction performance by the population-level response function from the training-set. Thus, well-predicted cells appear toward the upper left. (B) PCA of the test-cell response functions. Each point represents one cell, summarizing variability in the response patterns. (C) Mean parallel-to-wound response functions for the 20 cells ($\sim 7\%$ of the test set) whose accelerations were predicted most accurately by the training-set population-level model. (D) Mean parallel-to-wound response functions for the 20 cells with the poorest prediction accuracy.

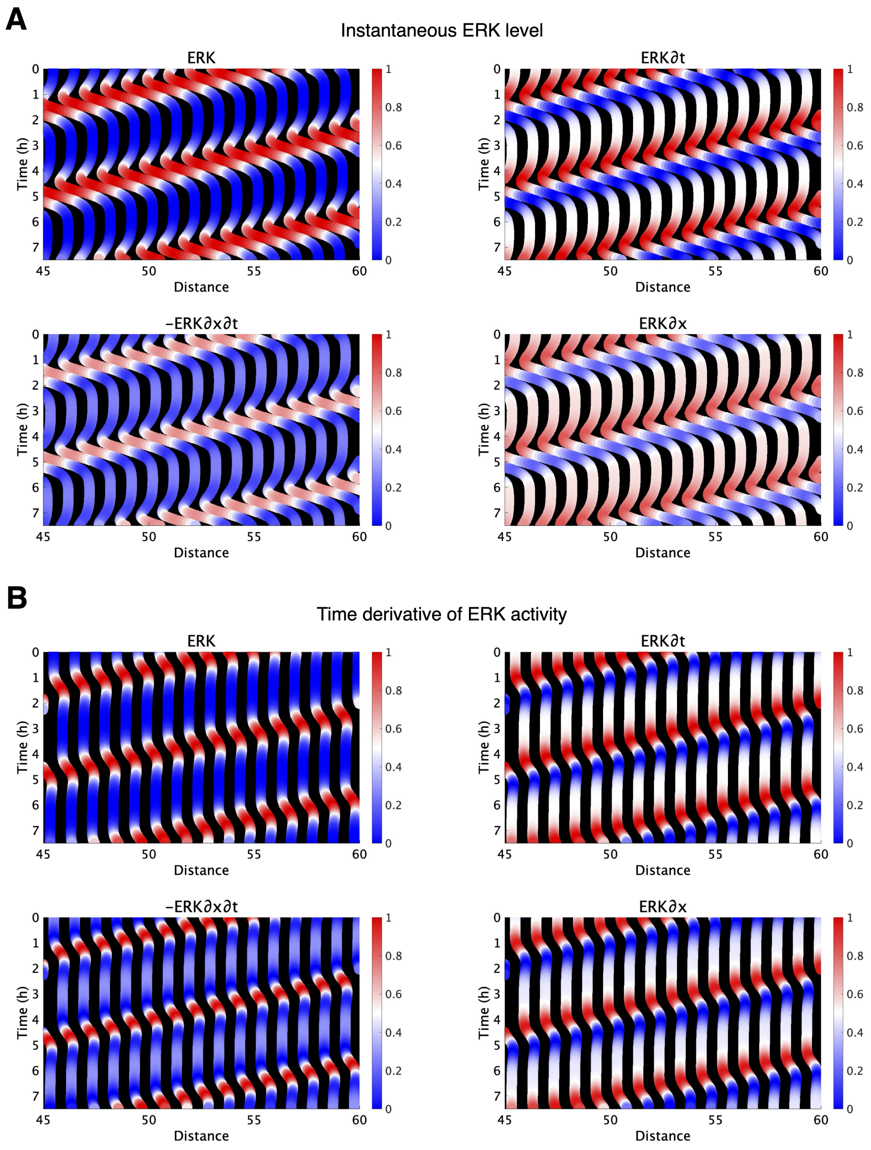


**Supplementary Fig S5** Numerical simulation of one dimensional particle and spring model. Horizontal axis: cell scan position (Distance, 45-60); vertical axis: elapsed time since the start of the experiment (Time, 0-7 h). Colors indicate the magnitude of each quantity. Top left: Heat-map of ERK activity, illustrating a wave that propagates from right to left over time. Top right: Temporal derivative of ERK activity; warm colors denote wave initiation, whereas cool colors correspond to regions where the wave subsides. Bottom left: Spatial gradient of the temporal derivative, capturing heterogeneity in the driving forces at the leading and trailing edges of the ERK wave. Bottom right: Spatial derivative of ERK activity, highlighting the steepness of concentration changes at the wave front and rear and revealing local features such as wave width and shape. (A)The instantaneous ERK level(Eq. 3 in previous study) (B)The time derivative of ERK activity A comparison of the top-left panels shows that quantifying the rate of change in ERK activity mitigates the apparent forward-propagation artifact observed in the previous forward Analysis. Parameter and setting of (A) are k = 2 (min 2), $\mu_0$ = 10 (min 1), Ro = 1/2, $\alpha$ = 1.5, $\beta$ = 2.5, $\sigma$ = 0.1 (min 1), sweeping velocity of illumination = 0.1 (min 1), width of illuminated area = 30, number of cells = 100. Parameter and setting of (B) are k = 2 (min 2), $\mu_0$ = 10 (min 4), Ro = 1/2, $\alpha$ = 10, $\beta$ = 2.5, $\sigma$ = 0.1 (min 1), sweeping velocity of illumination = 0.1 (min 1), width of illuminated area = 30, number of cells = 100. Where $k$ is the spring constant between neighboring cells; $R_0$ and $\mu_0$ are the basal cell radius and viscosity, respectively, modulated by ERK activity with amplitudes $\alpha$(size) and $\beta$(viscosity); and $\sigma$is the ERK decay rate.

### Supplementary Movie

**Supplementary Movie S1.** Collective cell migration and ERK Activity in Wound Healing of MDCK Cells. MDCK cells were scratched on the right side, and started collective migration to wound edge. Red and blue indicate high and low ERK activity, respectively.

### Supplementary Text

**Supplementary Text S1.** Below we derive a two-dimensional continuum model from our previous particle model:

$$\frac{dv_i}{dt} = -\mu_i v_i - k(R_{i-1} - R_{i+1}) + k(x_{i+1} - 2x_i + x_{i-1}) + \eta(v_{i+1} - 2v_i + v_{i-1}) \quad \text{(S1)}$$

Note that the equation holds for x- and y-directions. This model assumes cell mobility and cell area on the substrate to depend on the ERK acitivity. First, the ERK activity promotes cell mobility, which is effectively represented by reduction in cell-substrate friction as:

$$\mu_i v_i = \mu_0(-\beta \cdot ERK_i)v_i \quad \text{(S2)}$$

where $\beta$ is the coefficient for ERK-dependent friction and $ERK_i$ is the ERK activity in the $i$-th cell. Second, the ERK activity increases cell radius $R$:

$$R_i = R_0(1 + \alpha ERK_i) \quad \text{(S3)}$$

We approximate $(R_{i-1} - R_{i+1})$ as

$$\begin{aligned} R_{i\pm1} &= R \pm \frac{\Delta}{2}\partial_x R + \mathcal{O}(\Delta^2) \\ &= R_0\left[1 + \alpha ERK \pm \frac{\alpha R_0}{2}\partial_x ERK\right] + \cdots \end{aligned}$$

then

$$\begin{aligned} R_{i+1} - R_{i-1} &\simeq \alpha R_0 \Delta \partial_x ERK \\ &= \frac{\alpha R_0}{\rho}\partial_x ERK \quad \text{(S4)} \end{aligned}$$

For $(x_{i+1} - 2x_i + x_{i-1})$ ,

$$\begin{aligned} x_{i+1} - 2x_i + x_{i-1} &= \Delta^2 \frac{\partial^2 x}{\partial i^2} \\ &= \frac{1}{\rho^2}\partial_{xx}x - \frac{1}{\rho^3}\partial_x \rho \partial_x x + \mathcal{O}(\Delta^3) \\ &\approx -\frac{1}{\rho^3}\partial_x \rho \quad \text{(S5)} \end{aligned}$$

For the tissue viscosity term,

$$\begin{aligned}
\eta(v_{i+1} - 2v_i + v_{i-1}) &= \eta\Delta^2 \frac{\partial^2 v}{\partial i^2} + \mathcal{O}(\Delta^3) \\
&= \eta\left(\frac{1}{\rho^2}\partial_{xx}v_x - \frac{1}{\rho^3}\partial_x\rho\partial_x v_x\right) + \mathcal{O}(\Delta^3) \\
&\approx \eta\left(\frac{1}{\rho^2}\partial_{xx}v_x - \frac{1}{\rho^3}\partial_x\rho\partial_x v_x\right)
\end{aligned}$$

In our previous model, the tissue viscosity for $v_x$ depended only on velocity gradients in the x-direction; however, we expect a comprehensive constitutive formulation to include viscous stresses arising from velocity gradients in the y-direction as well. Therefore, the tissue viscosity term becomes

$$\frac{\eta}{\rho^2}(\partial_{xx}v_x + \partial_{yy}v_x) - \frac{\eta}{\rho^3}(\partial_x\rho\partial_x v_x + \partial_y\rho\partial_y v_y) \tag{S6}$$

Finally, expressing the time derivative of particle location in Eulerian coordinates gives

$$\frac{d}{dt} \to \frac{D}{Dt} \equiv \partial_t + \mathbf{v}\cdot\nabla \tag{S7}$$

Taken together(S1 - S7), we obtain:

$$\frac{Dv_x}{Dt} = -\mu_0(-\beta ERK)v - \frac{2\alpha k R_0}{\rho}\partial_x ERK - \frac{k}{\rho^3}\partial_x\rho + \frac{\mu}{\rho^2}(\partial_{xx}v_x + \partial_{yy}v_x) - \frac{\eta}{\rho^3}(\partial_x\rho\partial_x v_x + \partial_y\rho\partial_y v_y)$$

An analogous expression holds for by swapping $x$ and $y$. In our regression analysis, we omit the final term for simplicity because (i) the velocity Laplacian alone can represent tissue viscosity, and (ii) retaining the final term could lead to overfitting and induce numerical stability. Thus, we obtain Equation 2 in the main text.

$$\frac{Dv_x}{Dt} \simeq -\mu_0(1 - \beta ERK)v - \frac{2\alpha k R_0}{\rho}\partial_x ERK - \frac{k}{\rho^3}\partial_x\rho + \frac{\mu}{\rho^2}\partial_{xx}v_x + \frac{\mu}{\rho^2}\partial_{yy}v_x \tag{2}$$
